# Chemosensitive brainstem and diencephalic components in breath-hold fMRI

**DOI:** 10.1101/2025.04.11.648311

**Authors:** Simone Cauzzo, Alejandro Luis Callara, Maria Sole Morelli, Valentina Hartwig, Santa Sozzi, Francesca Frijia, Fabrizio Esposito, Marta Bianciardi, Domenico Montanaro, Claudio Passino, Michele Emdin, Alberto Giannoni, Nicola Vanello

## Abstract

**Objective:** The knowledge on breathing control and central chemoreception, key subcortical functions involved in several neuropathologies, is still mainly based on animal studies. In humans, functional MRI (fMRI) offers the needed spatio-temporal resolution and non-invasiveness, but the lack of specific tools and preprocessing solutions hinders its use in brainstem studies. We hereby propose an original fMRI analysis pipeline aimed at unravelling central chemoreception mechanisms, by integrating acquisition, spatial coregistration, noise removal and a novel data-driven analysis solution to compare network activation levels across tasks or conditions in fMRI.

**Approach:** Novel analysis methodologies are integrated with the optimization of known preprocessing approaches to physiological noise correction and brainstem-focused coregistration. We couple independent components of fMRI data, separately estimated from healthy subjects during Free Breathing (FB) and Breath Hold (BH), by means of spatial correlation. We then identify statistically significant differences between BH and FB in CO_2_-dependent components by means of voxel-wise comparisons of components’ percent signal change. Components were localized using the Brainstem Navigator Atlas for enhancing network interpretability.

**Main Results:** Using the pipeline we characterized CO_2_-related BOLD oscillations within the central control system of breathing. We corroborated the primary chemoreceptive role of medullary raphe in healthy subjects. We observed that BH over-activated ascending sensory-motor projections through the postero-lateral thalamus, descending projections through the putamen, and peripheral sensations entry points in the dorsal medulla. We highlighted the role of latero-dorsal tegmentum in the response to hypercapnia-induced aversive effects.

**Significance:** Our method allows to non-invasively locate primary chemoreception and related arousal triggers, characterizing alterations and therefore fostering the identification of therapeutic targets in abnormal breathing. Moreover, the proposed strategy addresses the issues of inter-task comparison among homologous independent sources, of their characterization and interpretation. Its extension could benefit all similarly challenging brainstem-focused studies, including those on Parkinson’s and Alzheimer’s diseases.

## INTRODUCTION

The central network of breathing control is a complex and heterogeneous system involving cortical, subcortical and brainstem structures that interact among them to form a closed circuit that regulates gas homeostasis in the blood [1], [2]. Non-invasive human studies focusing on this system could be crucial in the process of understanding and targeting central abnormal breathing patterns, such as central apneas or Cheyne Stokes Respiration (CSR) in Heart Failure [3], sudden infant death syndrome (SIDS) in preterm infants [4], or sudden unexplained death in epilepsy (SUDEP) [5]. Thanks to the favorable trade-off between spatial and temporal resolution and radiological non-invasiveness, fMRI is probably the most promising technique for accessing the small brainstem nuclei where central chemoreceptors and respiratory pattern generators are located. Recent investigations on the predictability of task-driven networks from resting-state studies increased the interest on the explorative value of fMRI, paving the way for a better integration between task and rest within the same study [6], [7]. Indeed, while spontaneous CO_2_ fluctuations are present at rest and constantly activate networks deputed to breathing control [8], gas challenge may amplify the neural response and strengthen the results, as well as enable to explore the system reaction to larger stimuli [9]–[11]. Nonetheless, exploring subcortical autonomic control networks also emphasizes the major weakness of fMRI, i.e., the low sensitivity and specificity of the BOLD signal in these regions. The hardest challenge in the study of respiration with fMRI is to disentangle specific BOLD fluctuations from non-specific effects, such as the vasodilatory properties of CO_2_, and the pulsatility of large blood vessels or CSF flow in large ventricles [12]. In addition, the small size of brainstem nuclei, the low temporal signal to noise ratio (tSNR) at the brainstem, and the presence of geometrical distortions and signal dropout in proximity of sinuses in the oral and nasal cavities further impair the study of the brainstem [13]. Ad hoc acquisition and pre-processing solutions tailored to the study of the brainstem have been recently proposed to overcome these problems [13]–[17]. Nonetheless, a comprehensive from-acquisition-to-analysis solution to the problem of closing the perduring gap between cortex and subcortex in MRI literature is still vacant, while most recent reviews still confirm how study design, registration and signal modelling in brainstem fMRI studies of the central autonomic network are open issues [18], [19]. Preliminary studies on healthy subjects based on standard model-driven approaches failed at describing CO_2_-related brainstem activations [20].

In this paper, we characterize brainstem, diencephalic and ganglionic CO_2_-related activity to unravel with fMRI the mechanisms supporting central chemoreception and breathing control in humans during Breath Hold (BH) and Free Breathing (FB). To this aim, we propose a robust and innovative dedicated pipeline that integrates both commonly used and custom approaches for the preprocessing steps, and a novel data-driven analysis solution for the comparison of network activation levels across tasks or conditions in fMRI. The pipeline is specifically dedicated to chemoreceptors’ mapping during BH, but it bears the power of being extended to autonomic unbalances in brainstem and diencephalic structures.

We defined an optimized strategy to deal with physiological noise and anatomo-functional image co-registration for the brainstem and the diencephalon, and the design of a novel inter-task comparison method. Physiological noise correction is based on the RETROICOR correction [21], which is regarded as a gold-standard physiological noise correction paradigm [22]–[26]. The standard approach is customized to improve the modelling of forced breathing patterns and compensate for the over-conservative approach in the study of the central control of breathing [20]. Specifically, based on the findings of this latter work, we here propose to optimize retrospective correction for the analysis of our target regions, by excluding respiratory-related regressors from standard RETROICOR.

Afterwards, image coregistration is guaranteed by means of is guaranteed by means of a customized multi-step procedure integrating partial oblique field of views with auxiliary opposite phase-direction encoded images, whole-head images and enhanced-tissue-contrast images.

To analyze the preprocessed data, we introduced a novel ad-hoc strategy based on masked Group Independent Component Analysis (mICA/GICA [27], [28]) and tested on BH and FB fMRI data. First, we highlight functional networks by matching independent components that share spatial patterns across conditions using inter-task spatial correlation. Second, we retrieve information on voxel-wise component activation strengths, to evaluate regional task-related modulation within each network. Particularly, we implement statistical tests between activation strengths expressed in percent signal change (PSC) [29]; we here propose to use this information, that is largely unexplored in the scientific literature, to quantitatively identify neural substrates within each network that are differentially modulated by FB and BH conditions. Third, we discuss findings integrating physio-anatomical information, including the most recent brainstem and subcortical atlases and available literature on the physiological control of breathing.

We provide critical advances to the knowledge of the human chemoreceptive and breathing control network, a key step in the direction of defining in detail the pathophysiology of abnormal breathing patterns, CSR, SIDS, or SUDEP. Moreover, we push the application of data-driven approaches to the exploration of complex, entangled dynamics in their evolution across sequential levels of task-based activation or pathology progression.

## MATERIALS AND METHODS

### Data acquisition

Ten healthy subjects (31±8 mean ± SD years old, balanced for sex) have been recruited. The experimental protocol was approved by the Institutional Ethical Committee, and it has been carried out in accordance with the World Medical Association Code of Ethics and the Declaration of Helsinki. All subjects signed a written informed consent. The absence of relevant pathologies or of MRI-incompatible prosthesis was assessed with a questionnaire. The acquired MRI data is not intended for public sharing due to privacy concerns. All analysis outcomes will be made available upon reasonable request to the author. Structural and functional MRI data were acquired using a 3 Tesla scanner (GE 3.0T HDx TWINSPEED, GE Healthcare, Waukesha, WI, USA). Structural MRI data were acquired using a three-dimensional T1-weighted sequence (3D FSPGR, TE=4.9, TR=10.7, FOV=25.6, voxel-size = 1 × 1 × 1 mm^3^). Functional MRI data were acquired using a T2*-sensitive gradient-echo echo-planar imaging sequence (TR = 2000 ms, TE = 30 ms, voxel-size = 3 × 3 × 3 mm^3^, Nr. of slices = 20, phase = 216). Partial coverage is motivated by the focus of this study on the brainstem, diencephalon (thalamus, subthalamus, hypothalamus) and basal ganglia (putamen, caudate nuclei, globus pallidum), and it was adopted to reduce SAR and maximize spatio-temporal resolution. Slices were parallel to the brainstem longitudinal axis and right-to-left phase encoding direction was adopted to minimize through-slice motion and blood-flow-related aliasing artifacts characterizing this region [14]. Acquisitions were repeated with the opposite phase encoding direction (left-to-right) to allow for distortion correction. No initial volumes were discarded in the acquisition phase during the pre-steady state of magnetization. Additional six-volumes acquisitions were performed on both phase-encoding directions with the same functional sequence but with whole brain coverage, to improve spatial coregistration of functional to anatomical images. Functional MRI data were acquired during 4 consecutive 7-minutes runs. As schematized in Fig. 1, during the 2nd and 4^th^ run, subjects performed a BH task, in which one minute of FB was followed by six cycles of 30 s of inspiratory BH alternated with 30 s of FB. Inspiratory BH (BH preceded by an inspiration) is reported to be easier to perform and to maintain for longer intervals, therefore minimizing motion and non-specific cognitive activity related to effort control [30]. During the 1st and 3rd runs, subjects were told to perform FB for the whole time. During the experiment, subjects had eyes closed and wore earplugs. For all runs, an operator standing in the MR room communicated to the subject the start and the end of each BH repetition by briefly touching the subject’s leg every 30 s. The same cueing was provided during FB to achieve a common baseline between conditions, but the subject was told to ignore these commands in FB runs. The right-to-left phase-encoding direction (polarity 1) was used for runs two and three, whereas the opposite phase encoding direction (polarity 2) was used for the other runs. This ensures the best configuration to extract for each run a nearby-acquired sub-sequence of 6 volumes having opposite phase-encoding direction and no BH challenge ongoing. The time-average of these 6 volumes was used to improve distortion correction (see Fig. 1). Concurrently, we acquired physiological signals: inspired-expired O_2_, expired CO_2_ and photo-plethysmography were recorded using a GE Datex-Ohmeda monitor, whereas breathing pattern was monitored using the MR-scanner pneumatic belt.

**Fig. 1.**
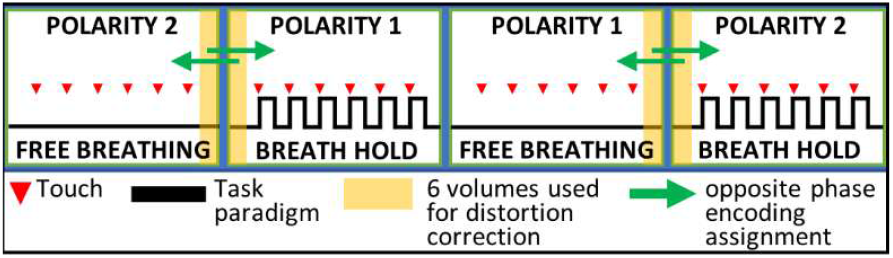
Acquisition sequence: Subjects performed a BH task in the 2nd and 4th runs (1’ FB, then 6 cycles of 30” BH + 30” FB). Subjects performed FB in the 1st and 3rd runs. Touches to the subjects’ leg (red arrows) are delivered every 30” and mark the beginning of apneas. Polarity 1: R-L phase-encoding direction.

### Data preprocessing

The preprocessing pipeline deviates from the most typical approaches to meet the following specific requirements: 1) the need for a cautious cleansing for physiological noise entangled with the signal of interest, through the customization of the RETROICOR approach; 2) the necessity to correct for larger distortions, overcome lower contrast-to-noise ratio and achieve enhanced coregistration precision in subcortical regions. Physiological signals recorded via the scanner-integrated pneumatic belt and the GE Datex-Ohmeda monitor were processed in Matlab (The MathWorks Inc., R2018b). End-tidal CO2 (PETCO2) was extracted as signal of interest from the envelope of expired CO2 maxima. Cardiac phase, and respiratory pattern peak timings and associated cycle amplitude were extracted to be used as input to retrospective correction of physiological confounds. A custom implementation of the standard retrospective physiological noise correction algorithm RETROICOR [21] was proposed. Specifically, more abrupt dynamics are allowed for the noise regressors related to breathing pattern, to cope with changes occurring during BH and right after the first inspiration. Moreover, these regressors are forced to constant values during the apneas. In addition, RVT regressors were discarded from the model. Particularly, we aimed at limiting the overconservative approach of the standard RETROICOR implementation, in the presence of forced or abnormal breathing patterns [20].

Anatomical images were skull-stripped and normalized to the MNI standard template with a nonlinear warp. Distortion-corrected whole head fMRI images were obtained by combining opposite phase polarity acquisition after standard preprocessing within AFNI (https://afni.nimh.nih.gov/). Then, the dataset consisting of 6 temporal images was split in two obtaining a base for inter-volume registration of partial-coverage EPI acquisitions, and a base for the coregistration with the anatomical image. The former was created by averaging the last four volumes, characterized by standard tissue contrast. The latter was created instead from the average of the first two volumes, characterized by enhanced tissue contrast due to the initial transient of magnetization. This coregistration base was first aligned to the skull-stripped anatomical image with an affine transform, then nonlinearly warped to it.

On partial-coverage fMRI images, spikes were removed from fMRI data, then the aforementioned custom implementation of RETROICOR was employed. After inter-slice temporal alignment, distortion correction was implemented by computing the median warp between the two 6-volumes portions of partial-coverage images selected as explained in Fig. 1. Intervolume rigid spatial registration was performed using as base the time-average of the last four volumes of cropped whole-head images. This base is already aligned to the enhanced-tissue contrast averaged first two volumes of the whole-head images, used as base for the coregistration with the anatomical image. We applied in only one step the transformation from the EPI images to the MNI standard template, along with a spatial resampling to a 2 mm isotropic grid: this transformation is composed of the distortion-correcting nonlinear warp, the rigid-body inter-volume registration, the affine registration and the nonlinear warp between the base for coregistration with the anatomical image and the anatomical image itself, and finally the affine and the nonlinear transforms between the anatomical image and the MNI standard template. The whole coregistration and normalization process is summarized in Fig. 2. On the spatially normalized data, spatial smoothing was applied with 3 mm FWHM gaussian kernel, and movement time series and their derivatives were regressed out.

**Fig. 2.**
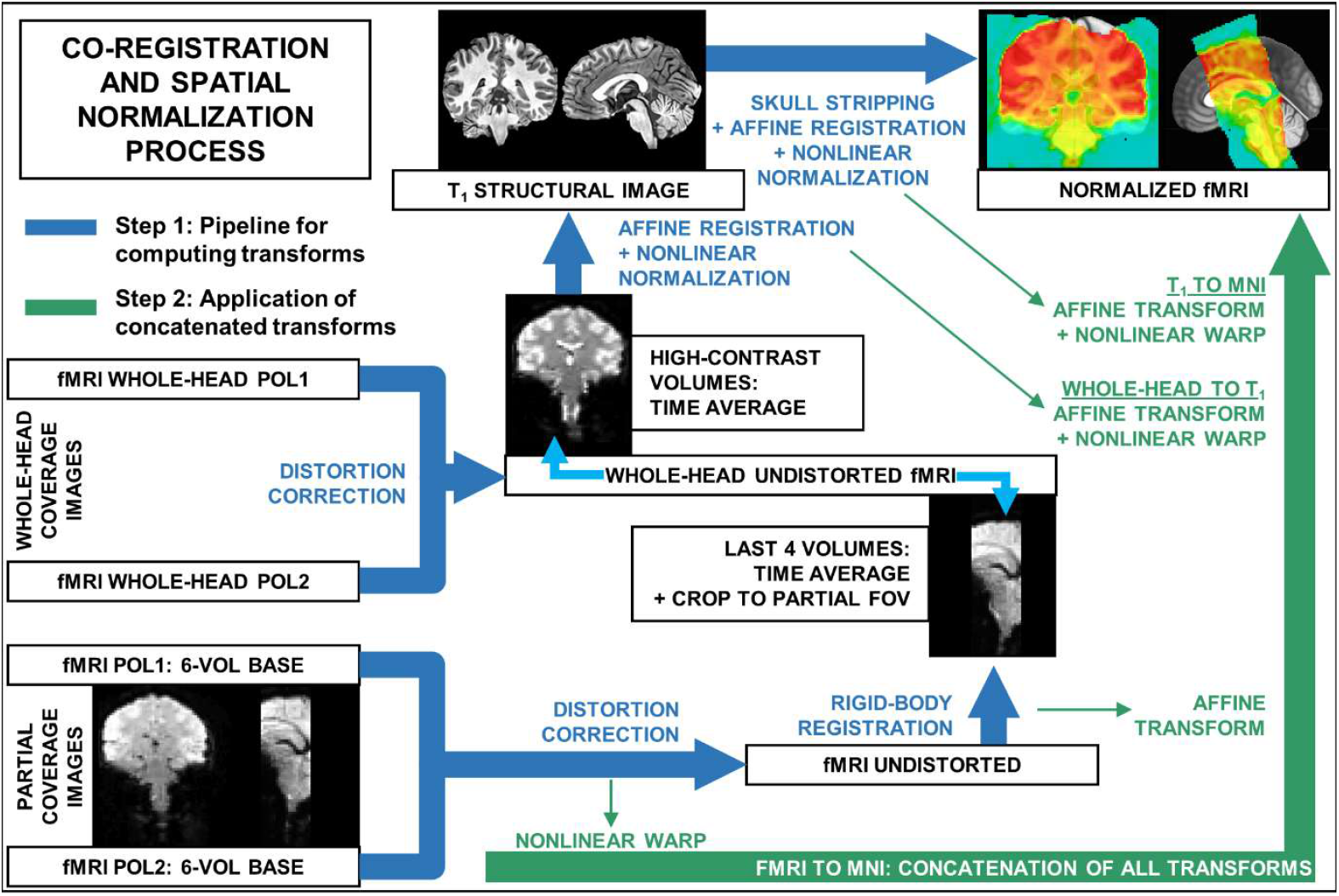
Coregistration-normalization process: the pipeline for computing single-step transforms (blue) and the process of their concatenation into a single transform (green). see Fig. 1 to visualize the extraction of partial-coverage 6-volumes bases for distortion correction. On the top-right, an example of a normalized partial-coverage functional image (semi-transparent heatmap) is overlayed on the MNI template (grayscale).

### Data analysis

For the core of the analysis, we employ two separate masked Independent Component Analysis (mICA, [27]) on the BH and on the FB group, in order to achieve three goals: 1) focusing on subcortical dynamics, avoiding the bias towards higher-SNR cortical ones [13], 2) employing a data-driven method to avoid analyses driven by hypotheses based on insufficient available knowledge on the neural response to CO2 oscillations; 3) designing a cross-task component comparison and characterization routine. On the results of these two analyses, we build a novel characterization strategy employing inter-task spatial association between different components and statistical comparison of activation levels.

First, a “subcortical mask” (Fig. 3A) was defined, comprising brainstem, diencephalon (thalamus, subthalamus, hypothalamus) and basal ganglia (putamen, caudate nuclei, globus pallidum). The mask is based on a parcellation provided with the OASIS-30 Atropos Template within the Mindboggle Project (https://mindboggle.info/data.html). The mask excludes the oro-nasal cavities and the ventricles anterior to the brainstem and in addition, it is eroded by two levels in the most anterior part of the pons and rostral medulla. This is motivated by the large signal dropout effects present in ventral brainstem regions, which cannot be resolved by distortion correction or precise coregistration and need to be masked out [31].

**Fig. 3.**
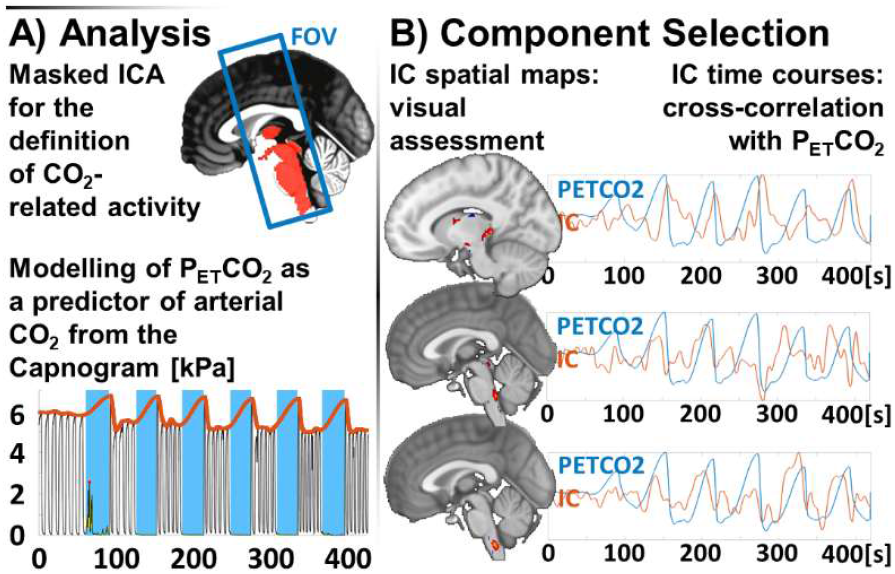
Data Analysis and Component selection. A) Masked Independent Component Analysis: top: the subcortical mask (red) within a blue square depicting the sagittal projection of the acquired field of view; bottom: example of PETCO2 signal (red) extracted from the envelope of expired CO2 partial pressure (black), with apneas highlighted in blue. B) Component Selection: three examples of independent components’ spatial maps and the respective time courses (red) aligned to express maximum correlation with PETCO2 (blue).

Two group-level mICA analyses have been performed, one on BH data and one on FB data, using the Matlab Toolbox GIFT ([32], http://icatb.sourceforge.net/). The GIFT Toolbox allows to estimate across-subjects components from concatenated data reduced with two-step subject-specific and group PCA. Nonetheless, it also provides subject-specific time courses and maps via back-reconstruction (GICA3 algorithm [29]). Time series were normalized for having spatial maps to express the PSC explained by the component, as allowed by GIFT calibration routines. This normalization allows to obtain parametric maps that can be compared between different runs or tasks, and it is widely used also in combination with model-driven approaches [33], [34]. Single subject ICA was implemented using the fastICA algorithm [35], [36]. We used the ICASSO algorithm, embedded in GIFT, to run ICA 100 times to obtain stable estimates [37]. ICA model order has been estimated to 29 components using Akaike Information Criterion (AIC), as implemented in GIFT.

### Component selection

Extracted components were selected by evaluating: 1) their spatial relationship with the underlying anatomy and 2) their temporal relationship with the PETCO2 time course (Fig. 3B). The spatial evaluation has been conducted via visual assessment of group-level spatial maps, converted to Z-scores and thresholded at |Z| ≥ 2, according to the criteria in [38], and before temporal correlation to avoid potential bias in the decision. Temporal evaluation was based on the cross-correlation of the subject-level time course with the associated pattern of PETCO2. Cross-correlation was computed considering lags in the [0-24] s range, to account both for circulation and pathophysiological delays between breathing-related CO2 changes and brain response [39]. A single maximum value was extracted for each component across lags. Accordingly, we obtained a total of 20 values (2 runs for 10 subjects) of maximum cross-correlation. Finally, a component was defined as PETCO2-related any time the cross-correlation significantly differed from zero as tested with a 2-sided t-test at p ≤ 0.05. Given the pseudo-constant dynamics of PETCO2 during FB, the selection based on temporal correlation was conducted only on BH components. All tests were corrected for multiple testing using a Benjamini-Yekuteli correction [40] as implemented in the fdr_bh Matlab Toolbox [41].

### Component characterization

After selection, components were further characterized through a three-step procedure based on their spatial similarity across tasks, the localization of the activation, and a voxel-by-voxel comparison of how homologous components are differentially modulated by the FB or BH. The discussion on their physiological meaning in the context of available literature will serve as further validation step.

#### 1) Spatial similarity across tasks

the stability of the activation pattern as described by a given independent component from BH data is evaluated by testing its similarity with maps of homologue independent components extracted from FB data, using Pearson’s correlation coefficients to associate each BH component with its most correlated FB one (in absolute value). The similarity is characterized in terms of four features: *BHvol*, expressing the volume of the BH |Z| ≥ 2 maps in terms of voxels; *ρ*, expressing the spatial correlation coefficient between the BH component and the associated FB one; *BH/OR*, expressing the overlap fraction between the BH |Z| ≥ 2 map and the logic OR between BH |Z| ≥ 2 and FB |Z| ≥ 2 maps; *overlap*, expressing the percentage of voxels of the BH |Z| ≥ 2 map that are included in the FB |Z| ≥ 2 map. The use of the absolute values in thresholding the maps for computing the overlap fraction mitigates possible effects of the sign ambiguity associated with the estimation of independent components [29].

#### 2) Localization of the activation

we computed the overlap between each independent map thresholded at |Z| ≥ 2 and the masks of regions of interest (ROIs) included in the recently-published Brainstem Navigator Atlas ([42]–[47], v0.9, https://www.nitrc.org/projects/brainstemnavig/), integrated with the Hypothalamus segmentation produced in [48] and the labels relative to Putamen, Thalamus, Caudate Nucleus and Pallidum from the parcellation produced in [49]. Brainstem Navigator labels cited in this work are laterodorsal tegmentum and central gray of the rhombencephalon (LDTG-CGPn), inferior medullary reticular formation (iMRt), medial geniculate nuclei (MG), parvicellular reticular nucleus alpha part (PCRtA), pontine nucleus oral and caudal part (PnO-PnC), raphe magnus (RMg), superior medullary reticular formation (sMRt), superior olivary complex (SOC), nucleus subcoeruleus (SubC), raphe obscurus (ROb), raphe pallidus (RPa), vestibular nuclei complex (VeC), viscerosensory motor complex (VSM). The components will be then named accordingly. In the brainstem, components overlapped with multiple close-by labels. Component location is expressed in terms of overlap fraction, reporting labels overlapping at least by 40% with the component |Z| ≥ 2 map.

#### 3) Activation strengths

PSC levels are compared between each couple of spatially similar components, obtained from FB and BH tasks respectively, using a t-test (p ≤ 0.05, FDR corrected using Benjamini-Hochberg), limited within the logical OR of the two |Z| ≥ 2 masks. This comparison is further characterized in terms of four indexes: *Tmax*, positive peak value for the t-statistic; *Tmin*, negative peak value for the t-statistic; *T+/OR*, fraction of voxels with significantly positive t-statistic for the BH-FB=0 test over the whole logic OR between BH |Z| ≥ 2 and FB |Z| ≥ 2 maps; *T-/OR*, fraction of voxels with significantly negative t-statistic for the BH-FB=0 test over the whole logic OR between BH |Z| ≥ 2 and FB |Z| ≥ 2 maps.

## RESULTS

### Component selection

In the BH group, 13 components were excluded at the spatial evaluation stage for being maximally intense within cerebral ventricles, white matter or tissue boundaries, or for reproducing one of the patterns described in Griffanti et al. [38] such as multiple small, sparse clusters with both positive and negative sign, or two close-by symmetric activations with opposite sign. Similarly, 15 FB components were excluded based on their spatial maps. 17 BH components were identified as PETCO2-related using a t-test on the maxima across lags of the cross-correlation functions (2-sided t-test at p ≤ 0.05, Benjamini-Yekuteli corrected). Seven of these components had already been discarded at the spatial evaluation stage of component selection. The remaining ten components were thus included in the final selection and considered for results characterization and discussion.

### Component characterization

Each BH component was associated with the most spatially similar FB component, as determined by Pearson’s correlation coefficient, resulting in 10 pairs of matched components, none involving discarded ones. The parametric maps and the features that we used to discuss matched components are shown in Fig. 4, Fig. 5 and Fig. 6. To evaluate the similarity between the two associated components, we report the BH component map thresholded at |Z| ≥ 2 (both as 2D slices and as a 3D rendering), a 3D rendering of the logic OR between the BH and the FB component thresholded at |Z| ≥ 2, and the values assumed for the features *BHvol, ρ, BH/OR* and *overlap*. The statistics describing the overlap can be found under the component’s name in Fig. 4, Fig. 5 and Fig. 6. The correlation coefficient provided an indication of similarity over the whole subcortical mask: it was above 0.40 for all components except for the Inferior Pontine Component and the Dorsal Ganglionic Component, which showed negative correlation values. The *overlap* index provided instead an indication of similarity between thresholded |Z| ≥ 2 masks: for four components (Dorsal Ganglionic, Ventral Pontine, Inferior Pontine and Superior Medullary) it was lower than 0.40.

**Fig. 4.**
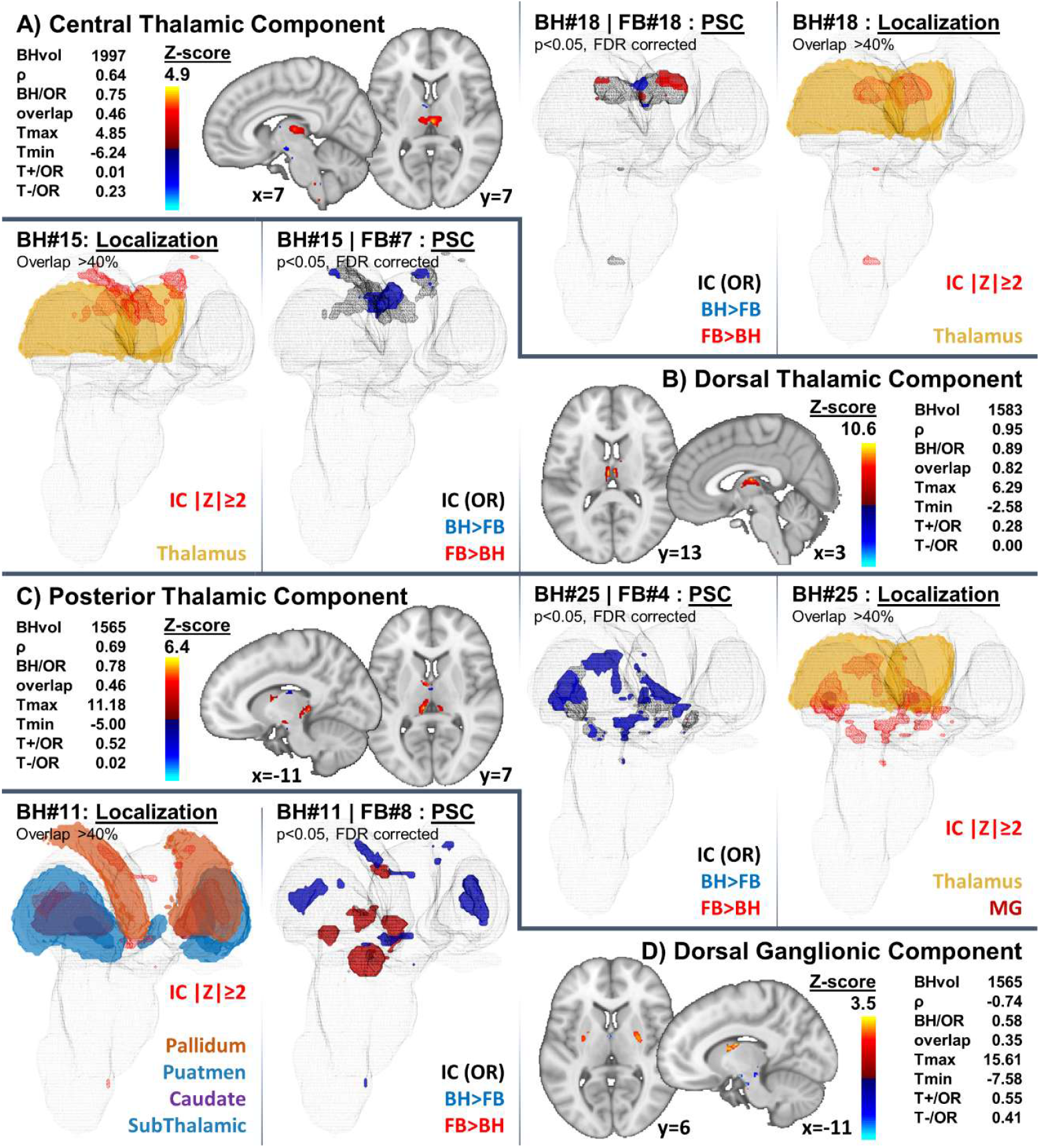
A) Central Thalamic Component, B) Dorsal Thalamic Component, C) Posterior Thalamic Component, D) Dorsal Ganglionic Component: for each component, we display: a sagittal and an axial 2D view of the Z-score map of the BH component, thresholded at |Z| ≥ 2; a 3D rendering for localization purpose of the thresholded Z-score map of the BH component (red mesh for both negative and positive clusters) and of the overlapping atlas labels (plain colors) within the subcortical mask (light gray mesh); the comparison between BH and FB in terms of PSC, displayed with a 3D rendering of the logic OR between the thresholded Z-score maps of the BH and FB components (black mesh) with the clusters of voxels showing statistical significance for the BH > FB condition in plain blue and those for the FB > BH condition in plain red (subcortical mask in light gray mesh). On one side, we report indexes characterizing the similarity between BH and FB components and the comparison between PSCs from both conditions.

**Fig. 5.**
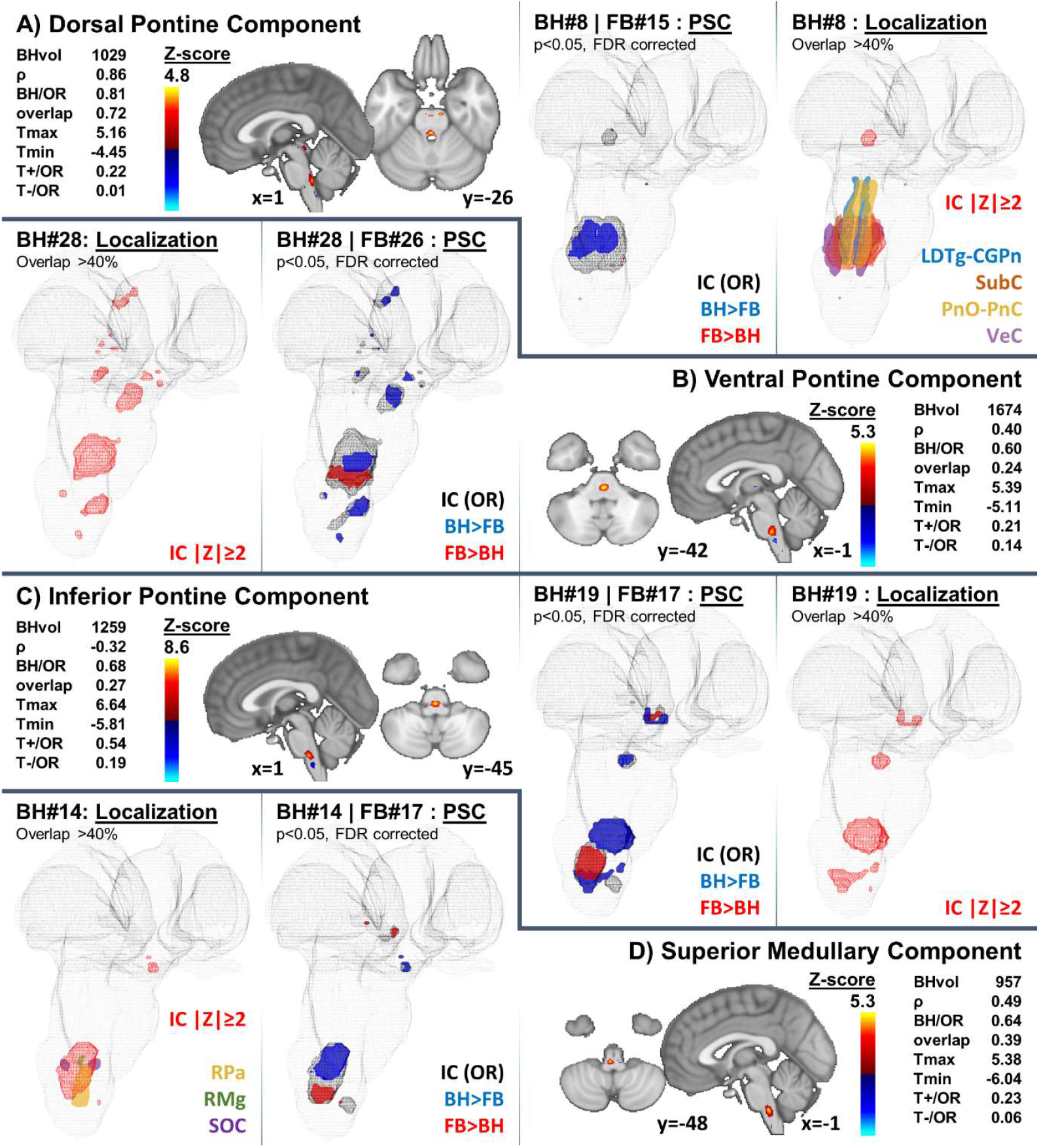
A) Dorsal Pontine Component, B) Ventral Pontine Component, C) Inferior Pontine Component, D) Superior Medullary Component: for each component, we display: a sagittal and an axial 2D view of the Z-score map of the BH component, thresholded at |Z| ≥ 2; a 3D rendering for localization purpose of the thresholded Z-score map of the BH component (red mesh for both negative and positive clusters) and of the overlapping atlas labels (plain colors) within the subcortical mask (light gray mesh); the comparison between BH and FB in terms of PSC, displayed with a 3D rendering of the logic OR between the thresholded Z-score maps of the BH and FB components (black mesh) with the clusters of voxels showing statistical significance for the BH > FB condition in plain blue and those for the FB > BH condition in plain red (subcortical mask in light gray mesh). On one side, we report indexes characterizing the similarity between BH and FB components and the comparison between PSC s from both conditions.

**Fig. 6.**
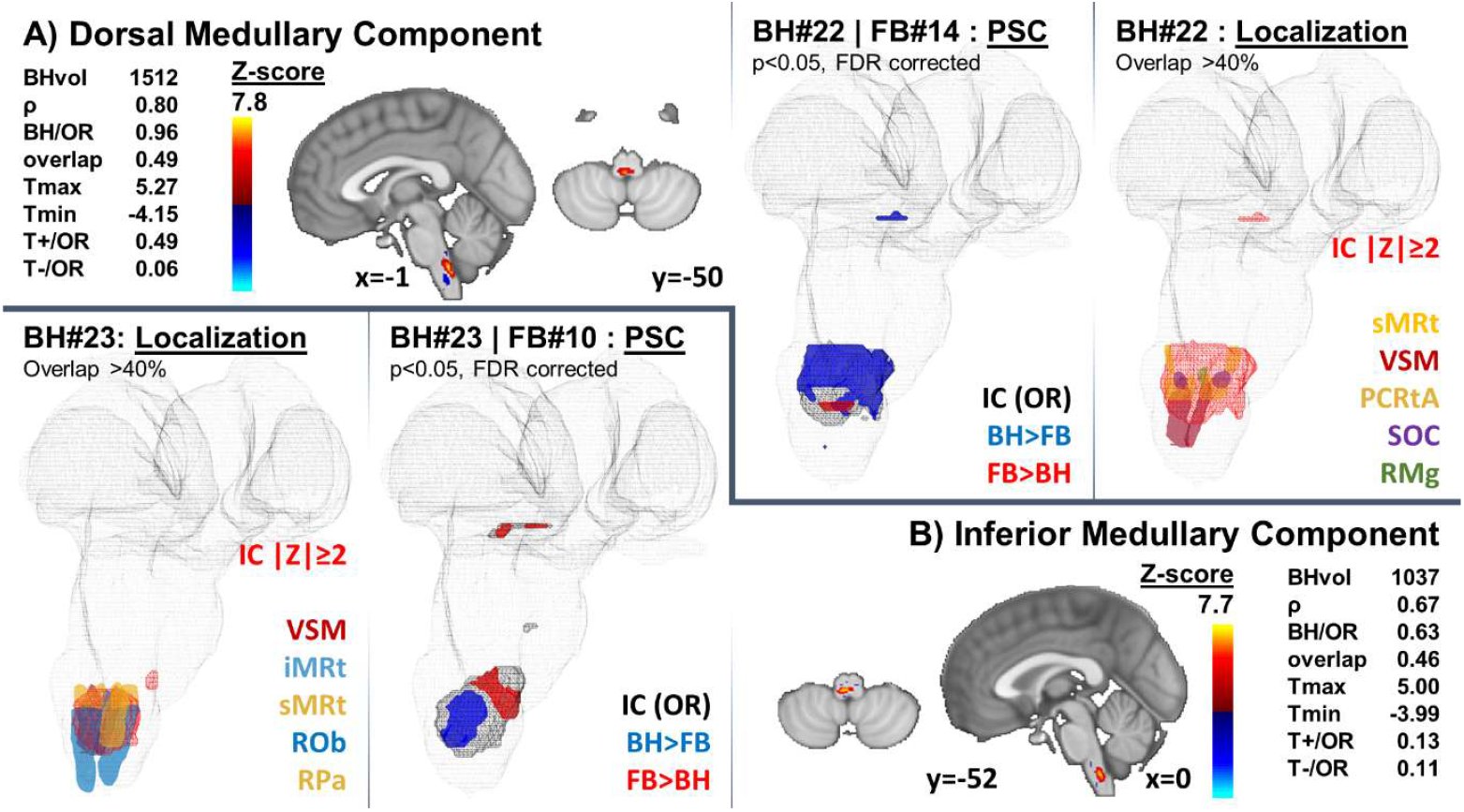
A) Dorsal Medullary Component, B) Inferior Medullary Component: for each component, we display: a sagittal and an axial 2D view of the Z-score map of the BH component, thresholded at |Z| ≥ 2; a 3D rendering for localization purpose of the thresholded Z-score map of the BH component (red mesh for both negative and positive clusters) and of the overlapping atlas labels (plain colors) within the subcortical mask (light gray mesh); the comparison between BH and FB in terms of PSC, displayed with a 3D rendering of the logic OR between the thresholded Z-score maps of the BH and FB components (black mesh) with the clusters of voxels showing statistical significance for the BH > FB condition in plain blue and those for the FB > BH condition in plain red (subcortical mask in light gray mesh). On one side, we report indexes characterizing the similarity between BH and FB components and the comparison between PSC s from both conditions.

The BH components were localized with the help of available atlases. Among the ten selected components, four showed an overlap above 40% with ROIs of the Brainstem Navigator Atlas, as assessed by evaluating the portion of voxels in the ROI mask overlapping with the component mask thresholded at |Z| ≥ 2. For the remaining components, two had the thresholded map entirely localized in the brainstem but did not meet the overlap threshold, and four were in basal or diencephalic regions. In the following, based on their location, we grouped the components in Diencephalic, Mesencephalic, or Medullary ones, and assigned new labels accordingly.

In the brainstem, components overlapped with multiple close-by labels. Considering 40% as threshold to assess the relevance of the overlap, we report in Table I the overlap fractions with the labels in the Brainstem Navigator Atlas for the four brainstem components that displayed at least one significant overlap. The Ventral Pontine Component (Fig. 5B) and the Inferior Pontine Component (Fig. 5C) were located above or in proximity of the ponto-bulbar junction, in the pons, also showing clusters in the mesencephalon: for them, no overlap above 40% was reported for any label of the Brainstem Navigator Atlas. In the Diencephalon, atlas ROIs were much bigger in size than those of the Brainstem Navigator Atlas. Therefore, localization was clearer and was left to a qualitative evaluation. The Central Thalamic Component (Fig. 4A) is entirely located within the two bilateral thalamic labels, whereas the Posterior Thalamic Component (Fig. 4C) showed a partial overlap with the thalamic labels and further extended over the medial geniculate bodies, covering their entirety; the Dorsal Thalamic Component (Fig. 4B) was located at the very top of the thalamic labels. The Dorsal Ganglionic Component (Fig. 4D) is entirely contained within the Putamen and the Caudate Nuclei labels.

**TABLE 1:**
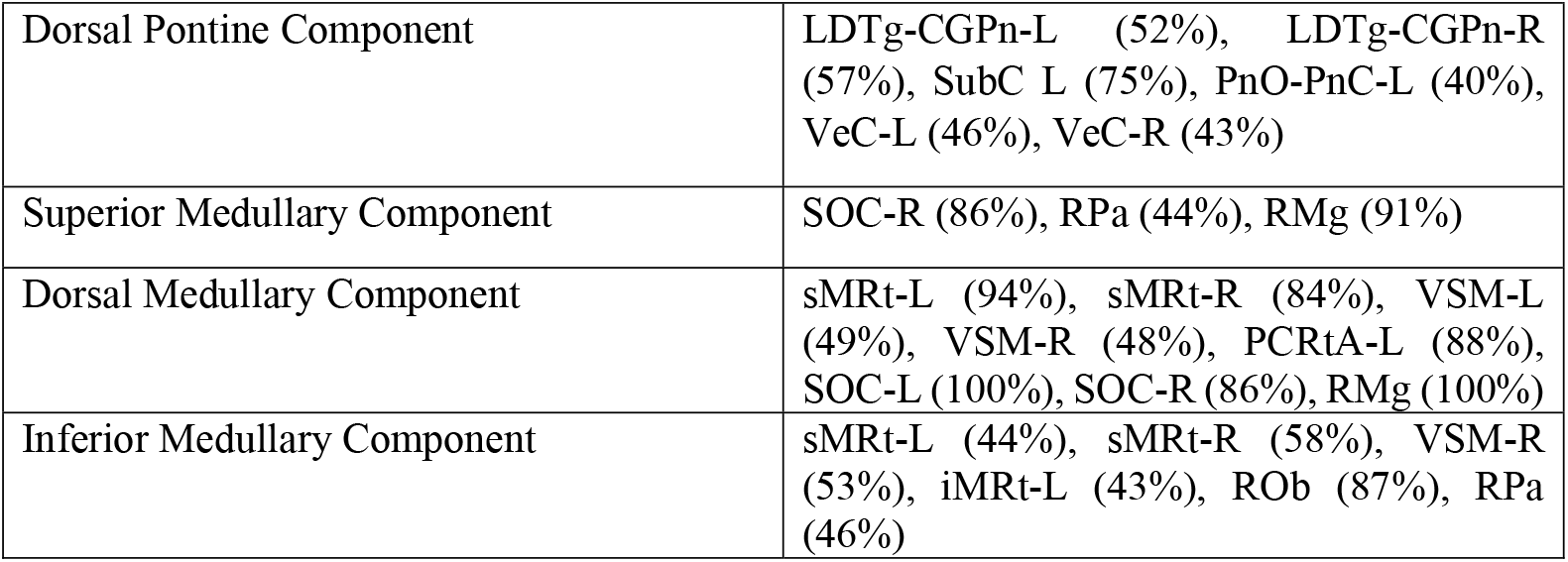
OVERLAP PERCENTAGES FOR BRAINSTEM NAVIGATOR LABELS OVERLAPPING BY AT LEAST 40% WITH BH COMPONENTS.

For the last component characterization step, activation strength, expressed as PSC, was compared voxel-by-voxel between matching BH and FB components, using a t-test. Results were visualized with 3D renderings in Fig. 4, Fig. 5, and Fig. 6, and reported in terms of *Tmax, Tmin, T+/OR* and *T-/OR* in (see below the name of each component). Two components, the Posterior Thalamic and the Dorsal Ganglionic, showed the highest *Tmax* (11.18 and 15.61 versus values between 4.85 and 6.29 for the other components), i.e., large maximal positive PSC deviations between BH and FB. *Tmin* values were more homogeneous across components. Components showing large values for *T+/OR*, i.e., widespread clusters of significantly higher PSC in BH with respect to FB, were the Posterior Thalamic, the Dorsal Ganglionic, the Superior Medullary and the Dorsal Medullary Components, all expressing values above 0.49, while the other components expressed *T+/OR* below 0.28. The Dorsal Ganglionic Component expressed large values for *T-/OR*, i.e., large clusters of significantly higher PSC in FB with respect to BH.

## DISCUSSIONS

In this work, we propose a novel framework to describe brainstem, diencephalic and ganglionic networks associated with breathing control during FB and BH tasks. The framework enables the inter-task association and comparison of group-level task-specific independent components. This is done in the context of a novel pipeline combining preprocessing and analysis and aimed at standardizing the investigation of subcortical autonomic processes, which are known to be highly entangled with oscillations in physiological dynamics such as blood gasses concentrations, heart rate or respiratory rate.

We highlight the relevance and the novelty of the implementation of our noise correction strategy. Physiological noise is among the major contributions to the BOLD signal fluctuations in highly perfused or CSF-surrounded brain regions, such as basal ganglia and cortical sulci. This is among the main reasons why we explored an approach based on masked ICA with a less aggressive correction for physiological noise. Based on previous work [20], we considered this solution particularly suited to avoid an overconservative approach and an erratic processing of physiological time series in the standard RETROICOR implementation, in the presence of abrupt breathing dynamics related to the breathing task. Indeed, in [20] authors focus on the analysis of both linear and nonlinear relationships of physiological signals and end-tidal CO2, within the context of BOLD signal change model in breath hold studies. Specifically, the description of BOLD signal change was improved using nonlinear functions of P_ET_CO_2_, and strong linear relationship was observed between RVT-related regressors and P_ET_CO_2_ changes. We here hypothesize, that RVT might still contain relevant information related to P_ET_CO_2_-based model and variability in circulatory delay [50].

As regards the proposed exploratory data analysis strategy, we believe that masked ICA plays a role in disentangling signals that could not be adequately separated with a GLM analysis. Particularly, imposing anatomical constraints to spatial ICA has been proven to, not only reduce the computational burden, but also improve the fitting of the ICA model in the grey matter voxels, in the case of fMRI data sets with a complex spatiotemporal structure [51]. The presence of a mild physiological noise correction step goes in the same direction of discarding sources of high variability with a conservative approach. The data-driven analysis can be thus focused on information that is not directly related to chest movements or cortical activity, sources that are prone to hide activity of interest. The physiological relevance of the findings described in this work is due to the integration of BH with FB data. The subcortical drive of breathing is based on a closed-loop system that modulates ventilation to maintain blood gasses concentration within physiological levels. The presence of subtle physiological oscillations of PaCO2 during resting state with a high impact on the BOLD signal motivates the presence of active networks deputed to breathing control both during FB and during BH [52]. In addition, several studies have recently focused on the ability of predicting task-related networks from resting state ones [6], establishing a parallel between the two [7]. Here, based on these studies, we hypothesize that it is possible to retrieve homologous components across analyses performed with ICA on BH and FB data, and that thereafter, it is possible to compare activation levels. Homologous components are defined in terms of spatial correlation between maps, considering the whole pattern without any threshold. Based on both the quantitative and qualitative (visual inspection of spatial maps) results reported in the previous section, correlation coefficients around ρ = 0.40 seem to provide a good clue for spatial correspondence, while between ρ = 0.30 and ρ = 0.40 the correspondence is debatable, and below ρ = 0.30 is indicatively absent. Assessing the quality of the spatial association across independent components is crucial, as the significance of the cross-task test on t-statistics depends on it. Once homologous components are recognized, percent signal changes are compared only within the region where either the BH or the FB components are significant, i.e., where |Z| ≥ 2. This was done to limit the detection of significant differences within the regions where at least one component is highly activated and to avoid possible significant differences arising from low activated regions. It is worthwhile noting that the component characterization was possible also thanks to the estimation at single subject level of PSC within each task. We observed that the potential of this approach is largely underestimated in the scientific literature, and therefore largely underexplored. To our knowledge, only in [53] it was proposed to perform inter-subject comparison of the components, and then in [54] the exploitation of estimated PSC was limited to the design of a model of bold signal changes during a CO2 gas challenge task. Here, we used such an approach to address inter-task group level characterization of components, thus allowing to improve the interpretability of the results.

### Physiological meaning

Given the strong interactions between cognitive and autonomic networks within the closed-loop control system of breathing, we consider both effects related to voluntary breathing control and to hypercapnia and homeostasis control, as they might be difficult to discern if one effect drives and one is driven by CO_2_ levels. Probably, the best possibility to discern between the two is given by a discussion on the physiology at the base of the activation. Here we discuss the selected components and their quantitative and qualitative characterization in the context of the literature on voluntary and autonomic breathing control, to assess the validity of the results and of the proposed methodology.

#### 1) Diencephalic Components

Four pairs of BH-FB components were individuated in diencephalic regions. Although chemoreception is expected to occur at the level of brainstem and Hypothalamus, diencephalic structures crucially connect them to higher order processing areas for the maintenance of homeostasis, in the transmission of alarming signaling and in conscious states management [55]. Indeed, the presence of projections from respiratory brainstem nuclei to thalamic nuclei has been observed in animals [56], [57].

The Central Thalamic Component did not convey substantial information on increased or decreased activations (*T+/OR* and *T-/OR* sum to only 24%). Its location, parted from the subcortical mask edges, speaks in favor of a specific activation, yet unmodulated by CO2 levels. Similarly, *T+/OR* and *T-/OR* sum to only 28% for the Dorsal Thalamic Component, which is placed at the very edge of the subcortical mask and thus prone to be driven by edge effects and CSF pulsatility. More interestingly, the PSC for the Posterior Thalamic Component and the Dorsal Ganglionic Component increases in BH at least within half of their |Z| ≥ 2 clusters. The spatial pattern of the Posterior Thalamic Component is in line with activity in ventral posterolateral nuclei linked with motor and somatosensory processing [11], reported in humans during CO2 challenges but also in animals [58], [59] and in sleep-disordered breathing patients [60].

Within the Dorsal Ganglionic Component, the Putamen is well known to be involved in both motor and autonomic functions, with major inputs received from autonomic-regulating cortical structures (e.g., the insula) [61], and in the switch between automatic and voluntary motor control [62]. Here, the increased activity of the Dorsal Ganglionic Component during BH can be associated with an overshoot in the input from cortical regions deputed to autonomic sensing. The presence of high anti-correlation within the whole component map, yet no overlap between thresholded maps, speaks for a key role in the switch between autonomic and voluntary control, or as alarm trigger. We would expect in this case that the same subcortical dynamics could be controlled by different inputs (a mesencephalic one in FB, a diencephalic one in BH) depending on the presence of cortical awareness of an unbalanced state.

#### 2) Pontine components

The Dorsal Pontine Component overlapped with bilateral LDTg-CGPn and VeC, but also monolaterally with left SubC and left PnO-PnC. The weighted centroid falls within the right LDTg-CGPn label, in close proximity to the medial line, therefore, overlap with SubC and PnO-PnC appears to be spurious. LDTg is mostly known for being involved in REM sleep and state change [63], [64], nonetheless, its projections to Ventral Tegmental Area (VTA) and Nucleus Accumbens were reported as crucial in mediating arousal and motor response to aversive stimuli [64], [65] as hypercapnia can be possibly considered. CGPn is instead scarcely reported in literature, in association to sexual receptivity [66]. VeC is implied in the known vestibular-autonomic interactions [67], the discussion of which is outside the scope of this paper, and is therefore left to future, more dedicated studies. The overactivation of the Dorsal Pontine Component in BH lets us hypothesize a link to the triggering of arousal or evasion action as a result of hypercapnic aversive stimuli, as this would be in line with aforementioned LDTg functions.

The major cluster of the Ventral Pontine Component was predominantly pontine, with minor clusters in the ventro-rostral medulla and in the ventral mesencephalon. The pontine cluster was possibly located in the area covered by the Reticular Tegmental Nucleus, Pontine Nucleus and medial lemniscus, respectively labelled RtTg, Pn and ml in [68], structures either deputed to motor control or to ascending sensory and proprioceptive pathways [69], [70]. The involvement of the ventral pons in the voluntary control of breathing was assessed by studying the “locked-in syndrome”: a lesion in the ventral pons was indeed associated with the impossibility of voluntary breathing acts [71]. In addition, a very similar activation was highlighted in a previous BH study [72] and linked to the integration of signals from supra-brainstem structures. We therefore associate this component with a top-down mechanism of behavioral override on autonomic breathing control. Clusters of *T+/OR* and *T-/OR* cover small portions of the |Z| ≥ 2 mask, respectively 21% and 14%. We hypothesize that it could be possible to isolate regions of voluntary motor control from regions of autonomic motor control, with the former placed slightly anteriorly.

The main cluster of the Inferior Pontine Component was located more caudally, at the ponto-bulbar junction, still showing no significant overlap with Brainstem Navigator Labels. This component was centered around slice z=-47 in MNI coordinates. Although the component spanned the whole width of the junction, peaking at the medial line, and although it was displaced at least 4 mm ventrally with respect to the centroids estimated from [68], we remark that the RTN position in the atlas of [68] is the result of extrapolations from animal studies, with no further evidence on humans. Some relation of BH Inferior Pontine Component with RTN chemosensitivity is thus not to be excluded. RTN chemosensitivity has been described as mediated by 5-HT release from serotonergic raphe nuclei [73]. This component is hardly matched by similar components in FB data, in line with the presence of structures activated only by alarm-triggering CO2 levels, as it is believed to happen for RTN [73].

#### 3) Medullary components

The Superior Medullary Component is of particular interest since it consistently overlapped with the RMg label, with its weighted centroid falling within it. BH and FB components showed *ρ* = 0.49, still, *overlap* was limited to 39%, with the FB cluster being slightly shifted inferiorly. From the comparison on PSC, we observed a major cluster of significantly positive t-value in the most superior portion, corresponding to RMg and rostral RPa. A significantly negative t-value was instead clustered in correspondence of the most caudal portion of RPa. The entirety of medullary raphe neurons was thus enclosed in a single network, with different levels of within-network activity. 5-HT neurons in the RMg and RPa are the main candidates to primary, unmediated chemoreception. Other candidates, such as RTN, were relegated to a secondary role by recent animal studies [73]. CO2-stimulated 5HT neurons are located in RMg and only the most superior portion of RPa [74]. RPa is also involved in thermal, vasomotor and sudomotor regulation [75]. As BH challenges and hypercapnia are known triggers of vasodilation and sympathetic activation [76], we interpret our results in the view of a dual function of medullary raphe neurons. BH increases the chemosensitive response, while inhibiting other autonomic functions.

The Dorsal Medullary Component spread over RMg as well, nonetheless, the weighted centroid of its BH component is further from the centroid of RMg than how the weighted centroid of BH Superior Medullary Component is (euclidean distance in voxels 3.4 vs 2.8, displaced caudally). We relate the Dorsal Medullary Component to a drive by sensory-motor nuclei VSM and sMRt. There is also bilateral overlap with the SOC label, which is nonetheless a vestibular complex involved in visual and hearing systems [45]. An involvement of the superior dorsal medulla in the voluntary control of breathing has already been observed in [77], where it has been associated with the activation of NTS and Nucleus Ambiguus (comprised in the ventro-lateral, superior VSM label), and explained as either a corticomotor-related excitation or inhibition at the medullary respiratory centers, or a sensory input from the lungs and chest mechanoreceptors through cranial nerves IX and X. We tried to limit this last contribution by asking volunteers to perform inspiratory BH without forcing the last inspiration, but anticipation phenomena might lead subjects to unconsciously exceed the normal expansion of lungs. Nonetheless, NTS is also known as the entry-point of peripheral chemoreception, which is expected to be triggered by BH-related decreases in PaO_2_ [2], [78]. This dual role of NTS as entry point of both peripheral chemoreception and peripheral mechanoreception motivates the clear increase in explained PSC in BH with respect to FB for the Dorsal Medullary Component.

Finally, the Inferior Medullary Component was centered around the ROb label. ROb has been recognized to play a role in the modulation of breathing frequency and amplitude, yet no primary role in chemoreception [79]. The correlation of BH Component time courses with PETCO2 is probably indirect and caused by a correlation with the respiration pattern during BH. In the comparison of PSC, a significant increase along the medial portion of the |Z| ≥ 2 mask, exactly around ROb, suggested an enhanced activation during forced breathing.

### Limitations

The sample size of 10 subjects is low but in line with the generally low number of subjects included in fMRI studies under complex physiological conditions and the related difficulties in increasing this number [80]. Nonetheless, we acknowledge that the study of low SNR regions in the lower brainstem would certainly benefit from using larger samples.

The tSNR being lower in inferior brainstem regions with respect to diencephalic regions [12] might play a role in decreasing the detection sensitivity of components originating from medullary activity. The meaning of negative correlation is elusive, despite the well-known ambiguity in the definition of the sign for an independent component [81]. In GIFT the sign is estimated from the component skewness, still this might seldom lead to misassignments. We observed that negative correlation was found for two pairs of components, the Dorsal Ganglionic Component, and the Inferior Pontine Component. These were also components showing overlap close to zero between the two thresholded masks. While for the Inferior Pontine Component we might assume that the -0.32 correlation coefficient was too low to define a homologous component, the correlation coefficient of -0.74 for the Dorsal Ganglionic Component clearly bound it with a CO2-dependent breathing drive and was thus discussed as such.

While 3 mm isotropic resolution is common for standard clinical 3 Tesla scanners, it still represents a limitation for the coregistration of the brainstem and the localization of activations, demanding thus ad-hoc solutions. Indeed, the Brainstem Navigator Atlas is based on multi-modal MRI acquired with 7 Tesla research scanners. Nonetheless, its translatability to 3 Tesla images has been proven by publishing 3 Tesla functional and structural connectomes of its labels [82]–[85] as the use on standard 3 Tesla images is a required step for the validation of these tools. Image interpolation is commonly employed in fMRI as a mild smoothing of data, which is known to increase SNR. Here, image interpolation is implemented by the GIFT software to resample independent components spatial maps to the grid of the anatomical reference. This solution allows to estimate the overlap also for the tiniest nuclei which could have disappeared if the atlas was subsampled.

Sensory stimuli were delivered to the subjects in all runs, to command the start and end of apneas in BH, and to trigger similar baseline activity in FB. Still, in the discussion of highlighted activations we ignored these stimuli. While the presence of the operator allows for better visual assessment of the task performance, the modality for providing the stimulus might be regarded as quite unprecise. We underline that in such an intrinsically block-designed study, the precision of this sensory cueing has low impact on the expected outcome of the single breath hold repetition. In addition, spatial ICA decomposition is time insensitive. Two factors concurred in excluding a major role for sensory stimuli in activating the observed subcortical networks: the PETCO2-based temporal criterion for component selection, and the anatomical localization of the independent component discussed on its physiological meaning. Residual correlation between PETCO2 and touch-triggered activations might be present since these touches are time-locked with the task. Still, brief somato-sensory stimuli are not expected to trigger block-design or ramp-like activity. On the contrary, activations related to BH have been modelled in literature with block designs [20] or with long double gamma functions [86]. The activity of chemoreceptors has been described as supralinearly related with blood CO2 levels [11], [54], [59]. Both activity patterns are expected to be well distinct from the event-related design of somato-sensory stimuli (see time series in Fig. 3), i.e., correlation is not expected to be significant. On another side, these touches impede defining FB runs as “resting state” and possibly disrupt certain resting state networks. Nonetheless, spontaneous oscillations of CO2 are still to be expected and as such, activity within the central breathing control network during FB.

### Significance

Studying the brainstem with fMRI on a conventional 3 Tesla scanner is crucial for promoting clinical applications. In this work, we tested to our knowledge for the first time the use of the Brainstem Navigator Atlas on 3 Tesla images for the localization of BOLD activations. The availability of the atlas considerably improved the possibility to discuss in detail our results, and to elaborate new hypothesis based on them. We highlighted activity in key chemosensitive brainstem regions and nodes of the breathing-control network, which could prove useful in studies of central abnormal breathing patterns. The validity of the pipeline is reinforced by the accordance between results and literature on breathing control: we linked activations in the ventral pons with the voluntary motor control of breathing, activations in postero-lateral thalamus with motor and somatosensory processing, and activations in the putamen with autonomic sensing. Moreover, the specificity of the methodology allowed to produce additional observations still lacking on human subjects and previously hypothesized from animal studies: we described in hypercapnic challenges CO2-related activity located around the Raphe Magnus, the main candidate for primary chemoreception, and in proximity of the Nucleus Tractus Solitarius, entry point for peripheral mechano- and chemo-sensation. Additionally, we suggested a role for LDTg as a mediator of arousal and motor response to aversive stimuli from the observation of its over-activation in BH. The translation of this

methodology to data acquired from pathological subjects would allow the comparison of case groups with controls, or different levels of pathology progression within the same group. Specifically, regarding the respiration control network, the presented methodology could be applied to the study of Cheyne-Stokes respiration in Heart Failure patients [3], [87], [88]. Moreover, given the relevance of the brainstem for the early insult of neurological disorders, such as Parkinson’s and Alzheimer diseases [89]–[92], the proposed pipeline has the potential of increasing the analysis specificity and reveal new biomarkers. In fact, beyond the relevance of the presented results, the application here pursued can be seen as a proof of concept for an extension of the framework to a broader set of studies on the central autonomic nervous system. Indeed, most of these studies put the focus on brainstem, thalamic and diencephalic networks but are challenged by a strong entanglement between signal of interest and physiological noise, a low tissue contrast and low SNR, and a lack of hypotheses that can drive the analysis. Therefore, our data-driven solution can be exploited for a proper goal-oriented acquisition and preprocessing and adapted to compare different tasks or conditions to better characterize effects.

## CONCLUSION

In this work, we developed an original approach to the preprocessing and analysis of subcortical fMRI signals acquired during BH challenges. The proposed pipeline tackles open issues that are specific to the study of brainstem regions with fMRI, such as their precise coregistration with anatomical images, and the correction for physiological noise when it is most entangled with autonomic processes. In addition, we introduced a novel analysis strategy for the inter-task association and comparison of independent components, broadening the explorative capabilities of ICA. Validation of independent components is carried out by inter-task comparison of percent signal change values and by providing a discussion based on the available literature on the voluntary control of breathing and central chemoreception. Interpretation is produced using state-of-the-art atlases such as the Brainstem Navigator Atlas. We provide a broad view on the central network of breathing control, suggesting the possibility to confirm for the first time in humans a primary role in chemoreception for serotonergic neurons of the medullary raphe, the recruitment of dorsal medullary relay centers of peripheral sensation, and the presence of an ascending arousal pathway signaling aversive stimuli such as hypercapnia.

## Acknowledgment

Research partially funded by the European Union - Next Generation EU, in the context of The National Recovery and Resilience Plan, Investment 1.5 Ecosystems of Innovation, Project Tuscany Health Ecosystem (THE), Spoke 3 “Advanced technologies, methods, materials and heath analytics “CUP: I53C22000780001.

## Data and code availability statements

The acquired MRI data is not intended for public sharing due to privacy concerns. All analysis outcomes will be made available upon reasonable request to the author.

